# Structural basis of ATP-dependent high-fidelity epigenome maintenance

**DOI:** 10.1101/2021.10.01.462825

**Authors:** Juncheng Wang, Sandra Catania, Chongyuan Wang, M. Jason de la Cruz, Beiduo Rao, Hiten D. Madhani, Dinshaw J. Patel

**Affiliations:** Structural Biology Program, Memorial Sloan Kettering Cancer Center, New York, NY, 10065, USA; Department of Biochemistry and Biophysics, University of California, San Francisco, San Francisco, CA, 94158, USA; Chan-Zuckerberg Biohub, San Francisco, CA 94158, USA

**Keywords:** cryo-EM structure, DNA methylation, DNA cytosine-5 methyltransferase, SNF2 ATPase, ATP, DNMT5, *Cryptococcus neoformans*

## Abstract

Epigenetic evolution occurs over million-year timescales in *Cryptococcus neoformans* and is mediated by DNMT5, the first maintenance-type cytosine methyltransferase identified in the fungal or protist kingdoms. DNMT5 requires ATP and displays exquisite hemimethyl-DNA specificity. To understand these novel properties, we solved cryo-EM structures of *Cn*DNMT5 in three states. These studies reveal an elaborate allosteric cascade in which hemimethylated DNA first activates the SNF2 ATPase domain by a large rigid body rotation while the target cytosine partially flips out the DNA duplex. ATP binding then triggers a striking structural reconfiguration of the methyltransferase catalytic pocket that enables cofactor binding, completion of base-flipping, and catalysis. Unmethylated DNA binding fails to open cofactor pocket and subsequent ATP binding triggers its ejection to ensure fidelity. This chaperone-like, enzyme-remodeling role of the SNF2 domain illuminates how energy can be used to enable faithful epigenetic memory.

**Highlights:**

- Structures of DNMT5 reveal mechanism of ATP-dependent DNA methylation
- Hemimethylated CpG recognition triggers partial base flipping of the target cytosine
- Hemimethylated DNA induces rigid body rotation to activate the SNF2 ATPase domain
- MTase catalytic pocket is remodeled by the SNF2 ATPase to achieve specificity

## INTRODUCTION

Maintenance methylation of hemimethylated CpG sites produced by DNA replication is the key to faithful epigenetic inheritance of genomic methylation patterns (Hashimoto et al., 2008). The process is mediated by maintenance DNA methyltransferases (DNMTs) (DNMT1 in animals and MET1 in plants) that recognize hemimethylated DNA (hmDNA) produced by DNA replication. There is almost nothing known about such processes in the two other eukaryotic kingdoms, namely fungi and protista, which harbor millions of species. The fungal epigenetic inheritance of DNA methylation patterns was demonstrated in the 1990s in the ascomycete *Ascobolus immersus* (Rossignol and Faugeron, 1995), although the maintenance enzyme responsible were never identified. We recently demonstrated that DNMT5, a DNMT required for CpG methylation in the human pathogenic basidiomycete fungus *Cryptococcus neoformans* (Huff and Zilberman, 2014), is a maintenance enzyme, the first cytosine DNMT to be characterized biochemically in the kingdom fungi or protista (Catania et al., 2020; Dumesic et al., 2020). DNMT5 is required for full virulence of *C. neoformans*, indicating an important function (Liu et al., 2008) and making it a potential drug target to inhibit infection (Huff and Zilberman, 2014). Amazingly, *C. neoformans* does not harbor a *de novo* enzyme because the gene for the *de novo* enzyme was lost in an ancestor that lived at least 50 million years (Catania *et al*., 2020). Methylation is maintained by an epigenetic evolution mechanism which involves replication of methylation patterns by DNMT5, rare epimutations, and natural selection (Catania *et al*., 2020). Biochemically, DNMT5 is unusual in that it harbors a SNF2 ATPase domain and requires ATP for DNA methylation (Catania *et al*., 2020; Dumesic *et al*., 2020). Moreover, it displays absolute specificity for hmDNA *in vitro* (Dumesic *et al*., 2020), much greater than mammalian DNMT1, which shows considerable methyltransferase (MTase) activity on unmethylated DNA (umDNA) *in vitro* (Goyal et al., 2006) and even *in vivo* (Arand et al., 2012; Liang et al., 2002; Lorincz et al., 2002).

To understand mechanisms underlying the ATP dependence and exquisite specificity of DNMT5, we solved three high-resolution cryo-EM structures revealing DNMT5 in the apo state, the hmDNA-bound binary state, and a quaternary state involving hmDNA-bound DNMT5 in complex with the nonhydrolyzable ATP analog, AMP-PNP, and the S-adenosylmethionine (SAM) reaction product, S-adenosylhomocysteine (SAH). These structures complemented by biochemical assays reveal a remarkable cascade in which hmDNA recognition triggers partial base flipping of the target cytosine out of the DNA duplex, accompanied by a dramatic allosteric movement in the ATPase domain followed by nucleotide binding. The binding of ATP leads to catalytic loop insertion and the completion of substrate base flipping and restructuring of the catalytic pocket to enable cofactor binding. In addition, ATP binding also licenses ejection of non-specifically bound umDNA. Overall, these studies illuminate a fundamentally novel signal transduction role for a SNF2 ATPase in an allosteric cascade that supports the remarkable specificity of DNMT5 for hmDNA, a property that is likely critical for epigenome maintenance over million-year timescales.

## RESULTS

### Overall Structure of DNMT5

We expressed near full-length DNMT5 (Figure 1A) including all its structured domains (residues 58 to 2377) in insect cells. This truncated protein displays remarkable specificity for hmDNA like full-length DNMT5, with no detectable MTase activity on umDNA or on hmDNA in the absence of ATP (Figure 1B). To study the mechanistic basis underlying these two intriguing features, a 36-base-pair (bp) DNA containing a single hemimethylated CpG site (Figure 2A) was incubated with DNMT5 supplemented with the nonhydrolyzable ATP analog AMP-PNP, and SAM reaction product SAH. It was then subjected to gel filtration purification and used for single-particle cryo-EM studies.

**Figure 1.**
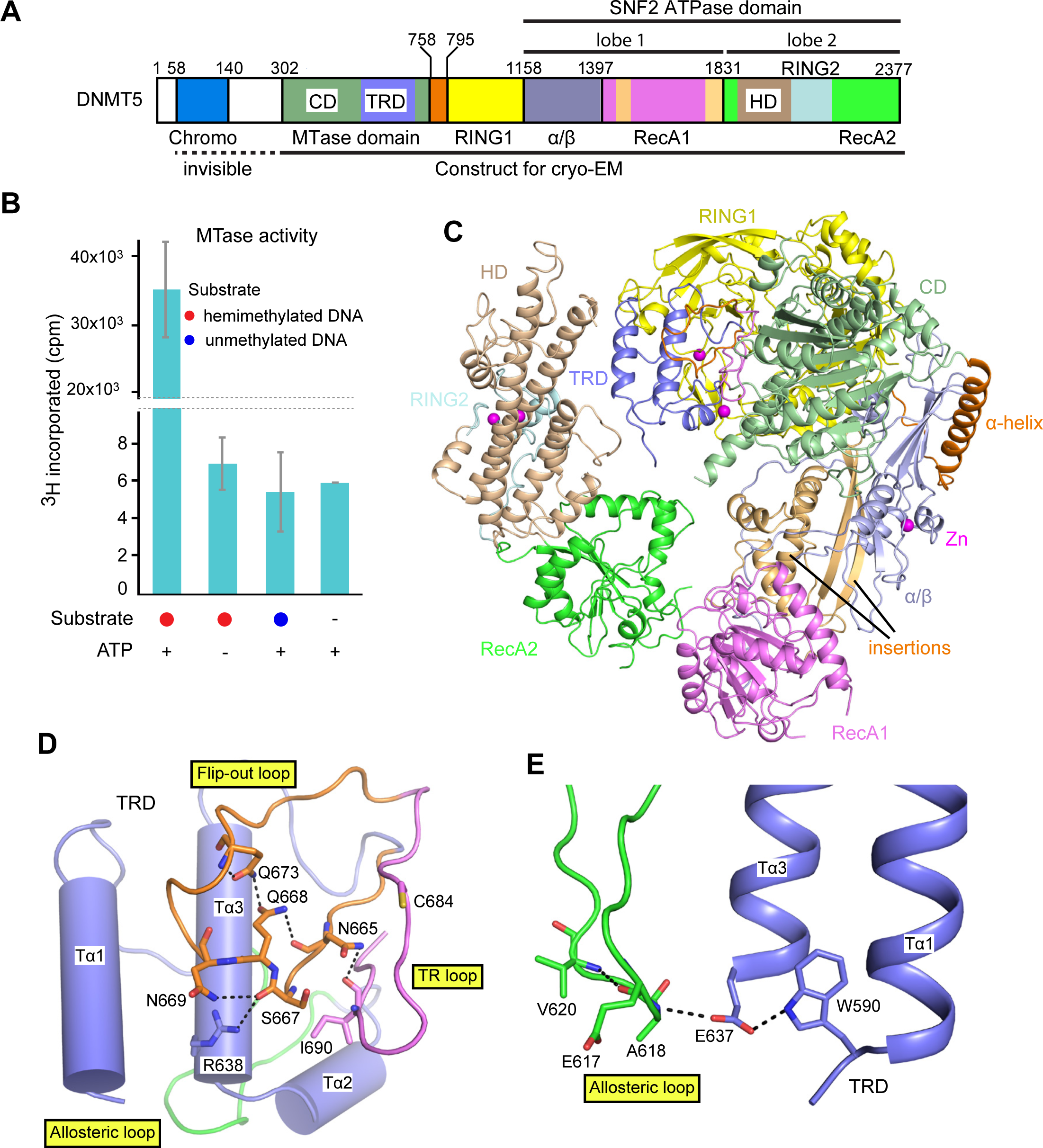
Cryo-EM structure of DNMT5 in the Apo State. (A) Color-coded domain architecture and numbering of DNMT5. (B) MTase activity of DNMT5 on umDNA and hmDNA substrates at the 4 hr time point. Error bars represent SD; n = 4. (C) Ribbon representation of the cryo-EM structure of DNMT5 in the apo state. (D) The relative alignments of flip-out and allosteric loops in the TRD of DNMT5. (E) A hydrogen bond network locks the allosteric loop through interaction with the TRD. See also Figures S1 and S2.

**Figure 2.**
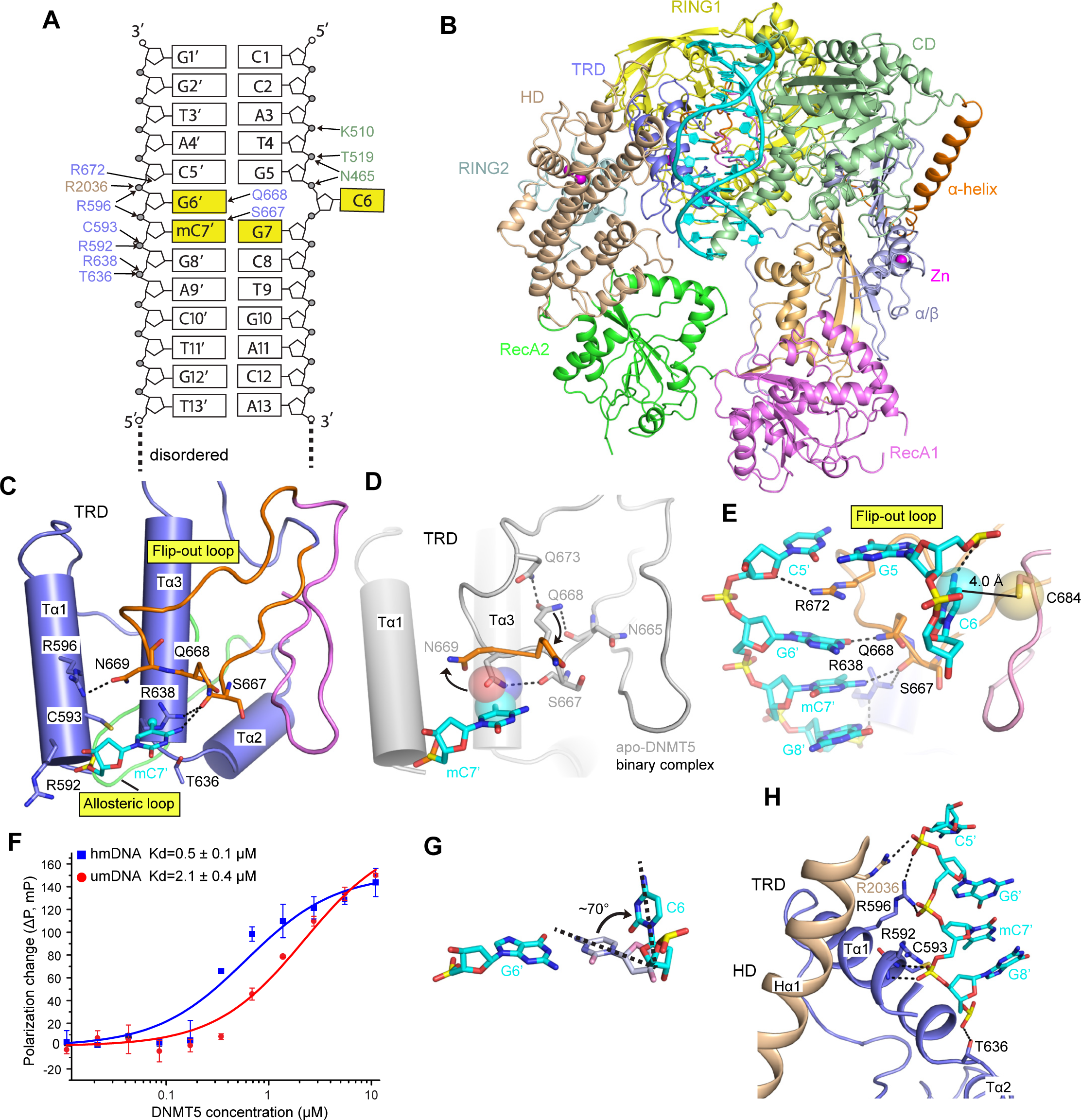
Cryo-EM Structure of DNMT5 Binary Complex with hmDNA. (A) The sequence of hmDNA including intermolecular contacts with DNMT5. (B) Ribbon representation of the cryo-EM structure of DNMT5 binary complex with hmDNA. (C) 5mC recognition by the TRD. (D) mC7’ induced conformational change of N669 and Q668 on binary complex formation. (E) Partial flip-out of the target cytosine C6 as a result of the interactions between the TRD and hmDNA. (F) Fluorescence polarization (FP) assay of FAM-labelled hmDNA and umDNA with increasing amount of DNMT5. Error bars represent SD; n = 3. (G) The partial flip-out of the target cytosine C6. (I) Interactions of DNMT5 with the phosphate backbone of hmDNA methylated strand. See also Figures S1, S2 and S3.

We obtained two structures through masked 3D classification with partial signal subtraction (Bai et al., 2015) (Figure. S1A), one of which is DNMT5 alone (apo-DNMT5) at 3.3 Å resolution (Figures 1C, S1B, S1C; Table S1) and a second composed of DNMT5 and hmDNA (binary complex) at 3.1 Å resolution (Figure 2B, S1B, S1D; Table S1). Side-chain densities of most residues in the binary complex map were clearly resolved, allowing accurate *de novo* model building (Figure S2A). The apo-DNMT5 structure was solved based on the binary complex model (Figure S2B). The N-terminal chromodomain lacks clear density in both the apo and binary complex maps, reflecting its flexible orientation relative to the other domains. Densities of AMP-PNP and SAH are absent in both maps, suggesting dissociation during purification.

DNMT5 folds into a compact structure with a ring-like topology (Figure 1C). The MTase domain is folded into two subdomains, labeled catalytic domain (CD) and target recognition domain (TRD). The TRD consists of three α-helices and two loops, designated as the flip-out and allosteric loops (Figure 1D). The sidechain of Q668 is stabilized by a hydrogen-bond network with Q673 and N665, while S667 interacts through hydrogen bonds with N669 and R638 (Figure 1D). On the back of the TRD, the allosteric loop is locked in place by two α-helices (Tα1 and Tα3) through a hydrogen-bonding network involving A618, E637 and W590 (Figure 1E).

A long rigid α-helix connects the CD and first RING zinc finger domain (RING1). The ATPase domain can be delineated into two lobes (lobe 1 and lobe 2). Lobe 1 contains an α/β region and the first RecA-like domain (RecA1) (Figure 1C). N- and C-terminal insertions of RecA1 make close contacts with the CD and contributes to the extension of the β-sheet in the α/β region (Figure S2C). Lobe 2 consists of the second RecA-like domain (RecA2) and an inserted region including an α-helical domain (HD) and the second RING zinc finger domain (RING2) (Figures 1A and 1C). The MTase domain, RING1 and lobe 1 compose the right segment, while lobe2 forms the left segment of DNMT5 (Figure 1C).

### Methylated CpG Dinucleotide Recognition

hmDNA binds within a cleft between the HD and CD, with the TRD accessing the DNA through major groove (Figure 2B). 13-bp of the 36-bp DNA encompassing the hemimethylated CpG site is visible in the binary complex map, implying the remaining flanking dsDNA is flexible and does not form stable contacts with the ATPase domain.

Sequence-specific intermolecular contacts between DNMT5 and hmDNA (Figure 2A) are made through the TRD. DNMT5 recognizes the 5mC (mC7’) by a hydrogen bond between the main-chain carbonyl of S667 and the N4 of mC7’ base (Figure 2C), which confers preference of 5mC over thymine. The 5-methyl group of mC7’ is accommodated in a shallow basic pocket within the TRD (Figures 2C and S3A). Such a 5mC accommodation is in marked contrast to the corresponding pocket in the structure of the DNMT1-hmDNA complex, where the methyl group of 5mC was anchored within a hydrophobic concave surface (Song et al., 2012). N669 from the flip-out loop also forms a hydrogen bond with R596 of Tα1 helix (Figure 2C).

Structural comparison of apo-DNMT5 and its binary complex reveals the side chain of N669 occupies the methyl binding pocket in apo-DNMT5 (Figures 2D and S3B). The binding of mC7’ shifts the side chain of N669 away from the pocket to interact with the Tα1 helix of TRD (Figure 2C). The conformational change of N669 in turn releases the side chain of Q668 from interactions with Q673 and N665 (Figure 2D), thereby positioning Q668 for recognition of the orphan guanine G6’ in the methylated CpG dinucleotide (Figure 2E). For the case of binding to umDNA, the side chain of N669 cannot be shifted away to initiate orphan guanine recognition. DNMT5 shows a 4-fold binding preference for hmDNA over umDNA of identical sequence (Figure 2F).

### A Novel Partial Flip-out of the Target Cytosine

The side chain of R672 penetrates deep into the DNA duplex forming a hydrogen bond with the deoxyribose sugar oxygen of C5’, with guanidinium group sandwiched by G6’ and C5’ bases (Figure 2E). The target cytosine (C6) partially flips out of the DNA duplex, oriented ∼70° from its pairing position, which is then stabilized by a hydrogen bond with the G5 phosphate (Figures 2E and 2G). This partially flipped-out state is strikingly different from the fully flipped-out state observed in other DNMTs or proteins and enzymes involved in writing, reading and erasing modified bases (Hong and Cheng, 2016; Klimasauskas et al., 1994). The partial base flipping does not require ATP as the target cytosine adopted the same partial flip-out conformation in the cryo-EM structure of DNMT5 solved by incubation with hmDNA and SAM but without AMP-PNP (designated pseudo-ternary complex) (Figure S3C). Notably, SAM density was not observed in the pseudo-ternary complex map.

C684 is in the proximity of the 5-carbon of the target cytosine pyrimidine ring (distance of 4.0 Å) (Figure 2E). It appears that C684 senses the methylation state of target cytosine, as a methyl group at 5-carbon would introduce steric hindrance. Notably, C684 is not the catalytic cysteine that initiates the methylation reaction.

DNMT5 makes sequence-specific contacts with the hmDNA only within the methylated CpG dinucleotide (Figure 2A) consistent with the lack of flanking sequence preference (Catania *et al*., 2020; Dumesic *et al*., 2020). hmDNA is further anchored by non-sequence-specific intermolecular interactions with DNMT5 (Figures 2H and S3D). R2036 from the HD makes direct contacts with hm DNA backbone.

### hmDNA Induces Profound Conformational Changes in DNMT5

Two views of the superposition of the structures of the apo-DNMT5 and the binary complex are shown in Figures 3A and 3B. The binding of hmDNA induces profound conformational changes in DNMT5, primarily observed within the left segment (Movie 1). The sequence-specific recognition by the flip-out loop (Figure 2E) facilitates the Tα1 and Tα3 helices of TRD moving closer to the DNA (Figure 3C). This movement disrupts the hydrogen-bond network locking the allosteric loop in the apo-DNMT5 structure (Figures 1E and S3E). The released allosteric loop contacts RING2, while RING1 also interacts with RING2 (Fig. 3D).

**Figure 3.**
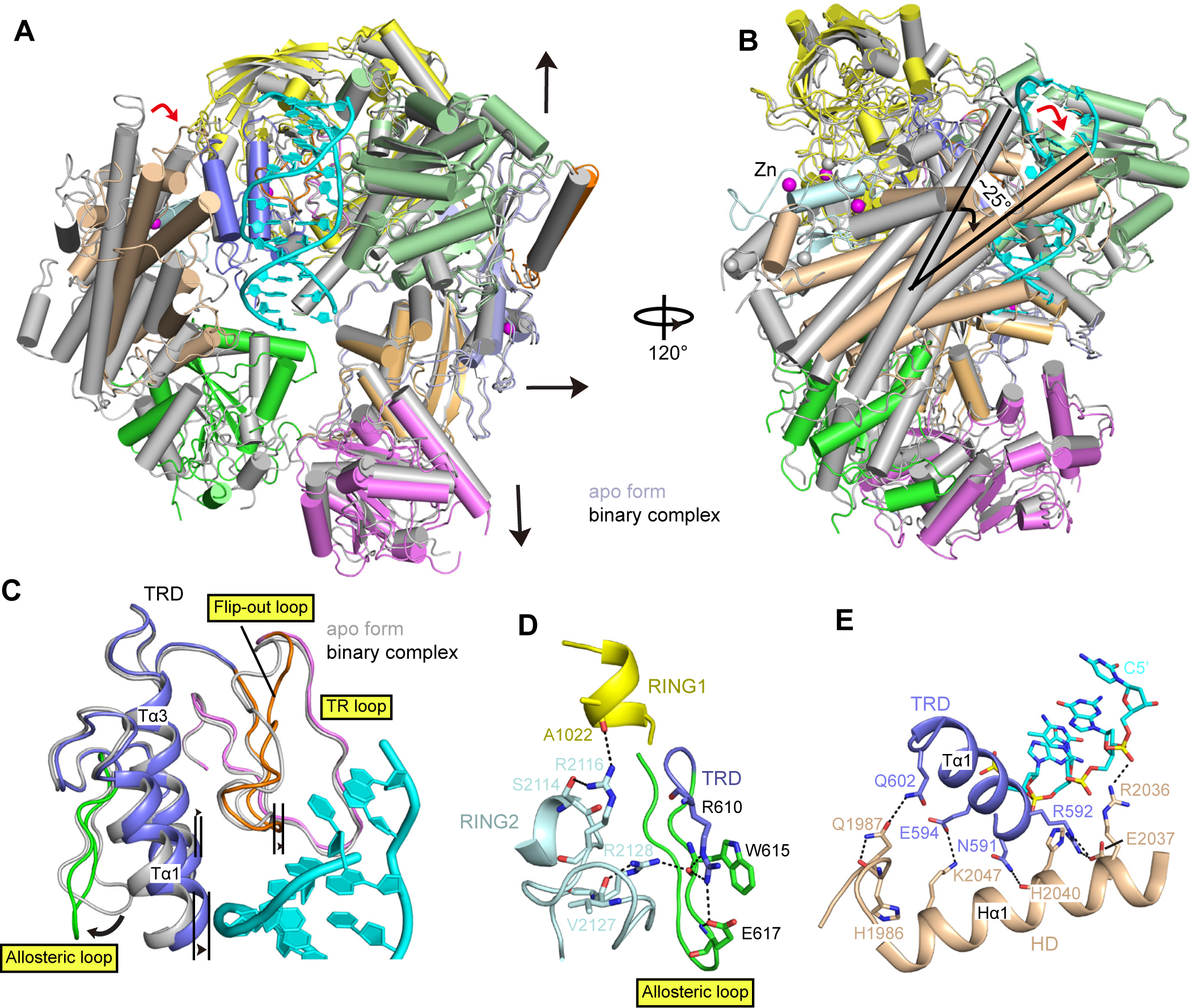
Conformational Changes in DNMT5 Induced by hm DNA Binding. (A and B) hmDNA binding induces conformational changes that expands the right segment (A) and rotates the left segment of DNMT5 (B). (C) hmDNA recognition by the TRD releases the allosteric loop. (D) Interactions between the allosteric loop and RING2, as well as between RING1 and RING2. (E) The hmDNA-induced conformational changes stabilized by interactions between the TRD and HD, as well as between the HD and hm DNA. See also Figure S3

As a result, the left segment of DNMT5 undergoes a substantial ∼25° clockwise rigid body rotation on binary complex formation (Figure 2B and S3F), with the two zinc atoms in the RING2 moving by 24.9 Å and 16.7 Å, respectively (Figure S3G). The rotated conformation is further stabilized by interactions between the TRD and HD, as well as between hmDNA and R2036 from the HD (Figure 3E).

hm DNA binding also impacts on the right segment of DNMT5, with the CD and RING1 moving upward, the α/β region moving towards the right and the RecA1 moving downward (see arrows in Figure 3A).

### hmDNA Binding Overcomes Autoinhibition in the SNF2 ATPase Domain

DNMT5 exhibits a basal ATPase activity of approximately 1 min^-1^, which can be stimulated ∼9-fold by hmDNA (Dumesic *et al*., 2020). The comparable binding affinity of DNMT5 to AMP-PNP without and with hmDNA (Figure S3H) cannot account for the stimulation. Structural analysis shows that the ATPase domain adopts an autoinhibitory conformation in apo-DNMT5 structure.

The Rα3 helix from RecA1 is in a face-to-face position with the Rα13 helix from RecA2 (Figure 4A) that would introduce a steric clash between the two RecA-like domains following formation of an ATP-induced closed conformation. The hmDNA-induced rigid body rotation moves the Rα13 helix away, thereby realigning the two RecA-like domains to allow domain closure for hydrolyzing ATP (Figure 4A). We mutated the Rα13 helix to a GGS loop (GS mutant) to relieve the autoinhibition. Indeed, the autoinhibitory effect was alleviated by ∼3.5-fold relative to wild-type (WT) DNMT5 in the absence of DNA (Figure 4B). The E637A mutant, which should facilitate release of the allosteric loop (Figure 1E), also increased the basal ATPase activity ∼2.5-fold (Figure 4B). These biochemical data support our “activation by rotation” model for ATPase activation.

**Figure 4.**
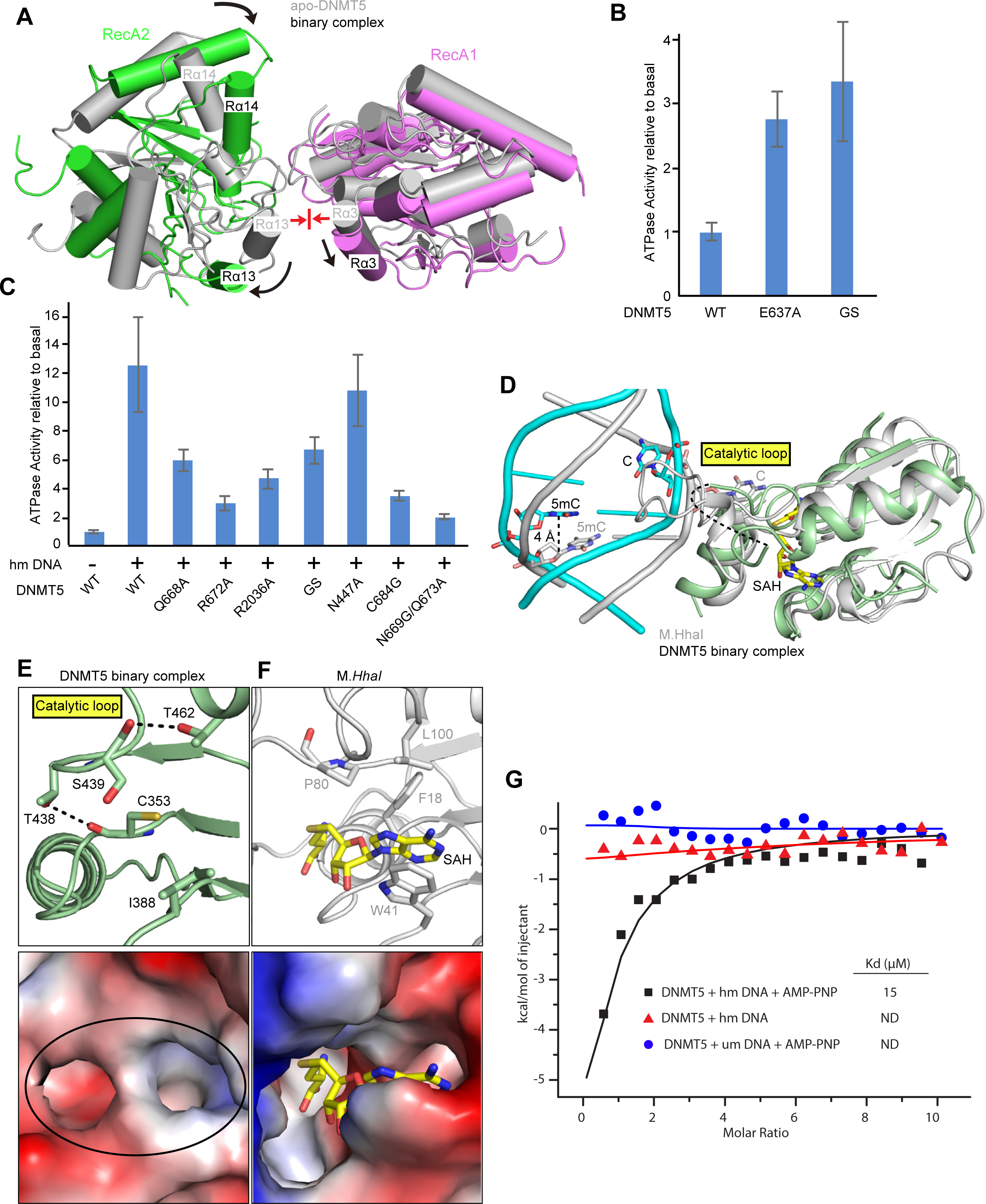
hmDNA Stimulates ATPase Activity while MTase Activity Remains Suppressed on Binary Complex Formation. (A) The realignment of the two RecA-like domains induced by hmDNA binding. (B and C) ATPase activity assay of wild-type (WT) and mutants of DNMT5 in the absence of DNA (B) and in the presence of hmDNA (C). Error bars represent SD; n = 4. (D) Structural comparation of DNMT5 binary complex with M.*HhaI* ternary complex with hmDNA and SAH (PDB ID: 5MHT). (E and F) Ribbon (top) and electrostatic surface (bottom) representation of the catalytic pockets in the DNMT5 binary complex (E) and in the M*.HhaI* ternary complex (PDB ID: 5MHT) (F). (G) ITC curves of SAH titrated into DNMT5 with indicated DNA or ligands. ND, not detectable. See also Figure S3.

We next mutated key residues involved in hmDNA recognition to test their impact on hmDNA-stimulated ATPase activity (Figure 4C). The ATPase activity of the N669G/Q673A double mutant, whose side chain cannot be shifted away to interact with Tα1 helix (Figure 2C), is least stimulated by hmDNA, even though the lock involving the Q668 side chain is weakened by the Q673A mutation. Disruption of the orphan guanine recognition through Q668A mutation (Figure 2E) reduces the ATPase activity ∼2-fold. Mutation of R672, that facilitates the partial flip-out of target cytosine through insertion into the duplex (Figure 2E), results in a 4-fold decrease of ATPase activity. The ATPase activity of R2036A mutant could not be effectively stimulated by DNA due to the disruption of the interaction between the HD and hmDNA backbone (Figure 2H). Replacement of the C684 that senses the methylation state of target cytosine by glycine (Figure 2E), reduces the ATPase activity more than 3-fold. These data are consistent with observations from the structural analysis.

### The MTase is Inactive in the Binary Complex

A DALI (Holm, 2020) search using the structure of the MTase domain in DNMT5 binary complex revealed *HhaI* methyltransferase (M*.HhaI*, PDB ID 3EEO) as the top hit (Z score 25.2). Structural alignment showed that the structure of DNMT5 MTase domain in the binary complex superposes well with M*.HhaI* ternary complex with hmDNA and SAH (O’Gara et al., 1996), with an r.m.s.d. of 3.0 Å for 140 Cα atoms (Figure 4D). However, the positions of hmDNA are strikingly different, with the 5mC-G step in DNMT5 binary complex positioned 4 Å above that in M*.HhaI* ternary complex (Figure 4D), such that the catalytic loop of DNMT5 cannot insert into the DNA duplex, with most of the loop invisible due to flexibility.

Notably, a small visible segment of the catalytic loop is stabilized by a hydrogen-bond network (Figure 4E). We compared the catalytic pocket of MTase domain in DNMT5 binary complex with its counterpart in M*.HhaI* ternary complex (Figures 4E and 4F). The side chains of the small visible segment of the catalytic loop in DNMT5 binary complex would introduce steric clashes with the bound SAH, thereby preventing cofactor binding. Indeed, no binding was detected when SAH was titrated into hmDNA pre-incubated DNMT5 without AMP-PNP monitored by isothermal titration calorimetry (ITC) (Figure 4G). This is also consistent with the absence of SAM density in the pseudo-ternary complex map. The majority of the catalytic loop is also invisible in apo-DNMT5 and the small visible catalytic loop segment acts like a lid that closes the catalytic pocket blocking SAM access to both apo-DNMT5 (Figure S3I) and its binary complex (Figure 4E). These findings provide a mechanistic explanation for no MTase activity for the binary complex in the absence of ATP (Figure 1B).

### AMP-PNP Binding Induces a Closed Conformation in the ATPase Domain

The limited binding preference for hmDNA and ATPase activity stimulation cannot explain the remarkable specificity of DNMT5. To further investigate molecular mechanisms underlying the remarkable specificity, we generated DNMT5 quaternary complex by incubation with hmDNA, AMP-PNP and SAH, followed by direct loading on the cryo-EM grid without further purification.

The quaternary complex structure was solved at 3.5 Å resolution with the densities for AMP-PNP and SAH readily traceable (Figures 5A and S4; Table S1).

**Figure 5.**
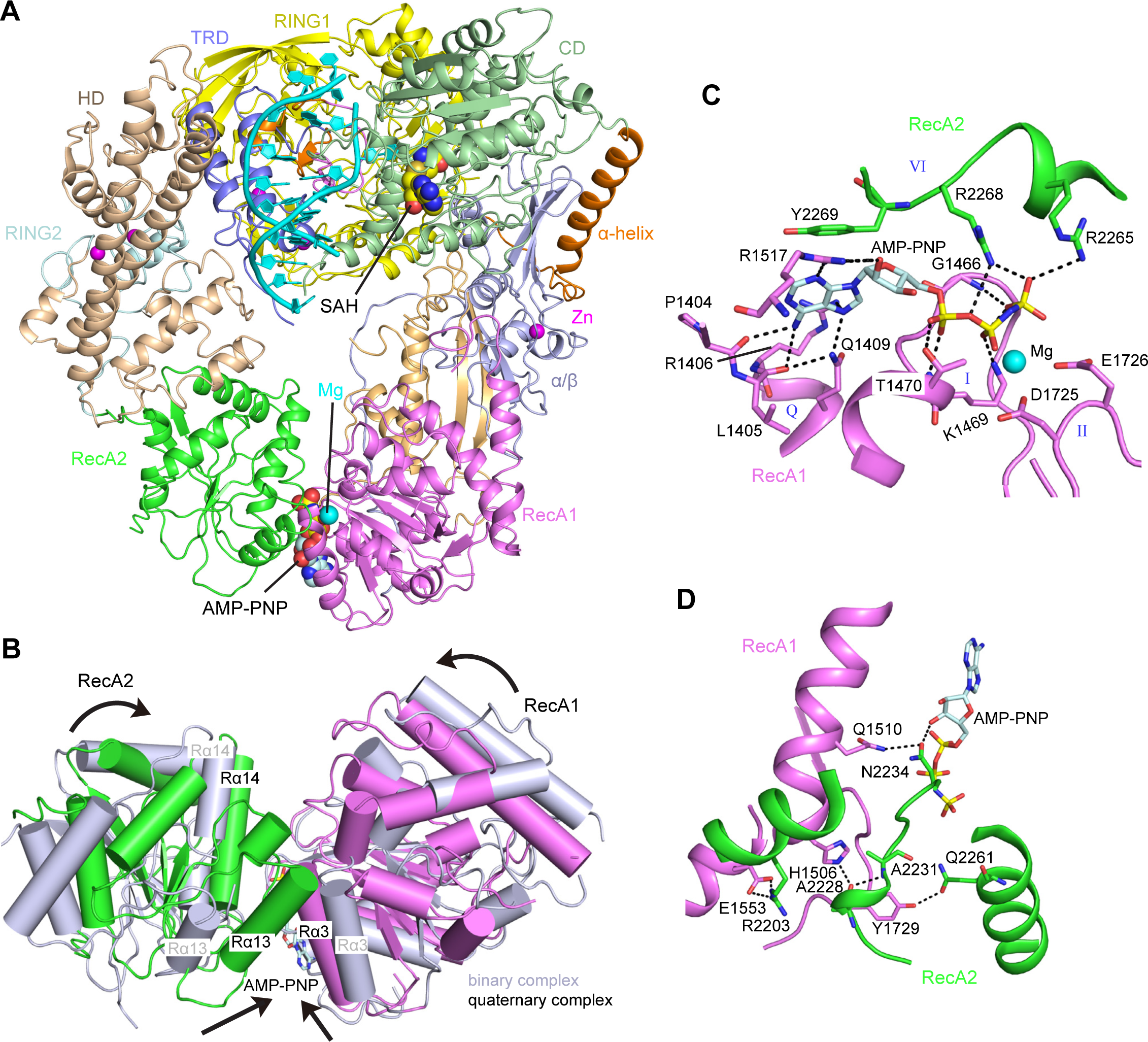
Cryo-EM Structure of DNMT5 Quaternary Complex. (A) Ribbon representation of the cryo-EM structure of DNMT5 quaternary complex with hmDNA, AMP-PNP and SAH. (B) The closed conformation of ATPase domain induced by AMP-PNP binding. (C) AMP-PNP mediated interactions with RecA1 and RecA2 domains. (D) Hydrogen bonding interactions between RecA1 and RecA2 domains. See also Figures S4 and S5.

The two RecA-like domains form a closed conformation upon AMP-PNP binding, with the cleft between the two domains closing around the bound ATP analog (Figure 5B). AMP-PNP binds primarily within RecA1, with residues from RecA2 contributing to recognition (Figure 5C). RecA1 contains the classic Q, I, II and III motifs, whereas RecA2 contains the IV, V and VI motifs (Fairman-Williams et al., 2010) (Figure S5). K1469 and T1470 of motif I are hydrogen-bonded to the α- and β-phosphates of AMP-PNP, while D1725 and E1726 of motif II coordinate the Mg^2+^ cation positioned between the β- and γ-phosphates (Figure 5C). R2268 and R2265 of motif VI are coordinateed to the γ-phosphate. In addition to the AMP-PNP-mediated interactions between the two RecA-like domains, residues from both domains also form direct hydrogen bond interactions (Figure 5D).

The adenosine ring of AMP-PNP is sandwiched through stacking with a tyrosine (Y2269) from motif VI in one direction and with the Q-motif, with which its forms intermolecular contacts, in the opposite direction (Figure 5C). A similar stacking platform sandwiching the adenine ring of cofactor is also observed in SF1 helicases, except that the aromatic residue is located in motif IIIa (Fairman-Williams *et al*., 2010). Such a two-directional sandwiching interaction appears to facilitate the ATPase domain to adopt a closed conformation more effectively in the ATP bound form. By contrast, a conserved tyrosine is positioned within the Q-motif of ISW1 and SWR1 (Figure S5), which are nucleosome remodelers of the SNF2 family, therefore enabling stacking from one direction.

We compared the ATPase domain of DNMT5 with SNF2 family members ISW1 and SWR1, whose structures have been reported in complex with nucleosome core particles (Willhoft et al., 2018; Yan et al., 2019). The key residues, locating within Ia-c, IV, IVa, V and Va motifs, that are required for DNA binding in chromatin remodelers (Fairman-Williams *et al*., 2010), are not conserved in DNMT5 (Figure S5). In addition, no density of interacting DNA was observed near the ATPase domain in the maps of both DNMT5 binary and quaternary complexes. Further, the isolated ATPase domain did not display DNA binding *in vitro* in our earlier study (Dumesic *et al*., 2020). These data suggest that the ATPase domain of DNMT5 does not harbor DNA binding capacity.

### SNF2 ATPase Domain Closure Allosterically Activates the MTase Domain

The closed conformation of ATPase domain formed upon AMP-PNP binding changes the relative orientation of RecA1 and RecA2 (Figure 5B). These changes are propagated to both segments of DNMT5, resulting in substantial conformational changes and an overall more compact structure (Figure 6A and Movie 2).

**Figure 6.**
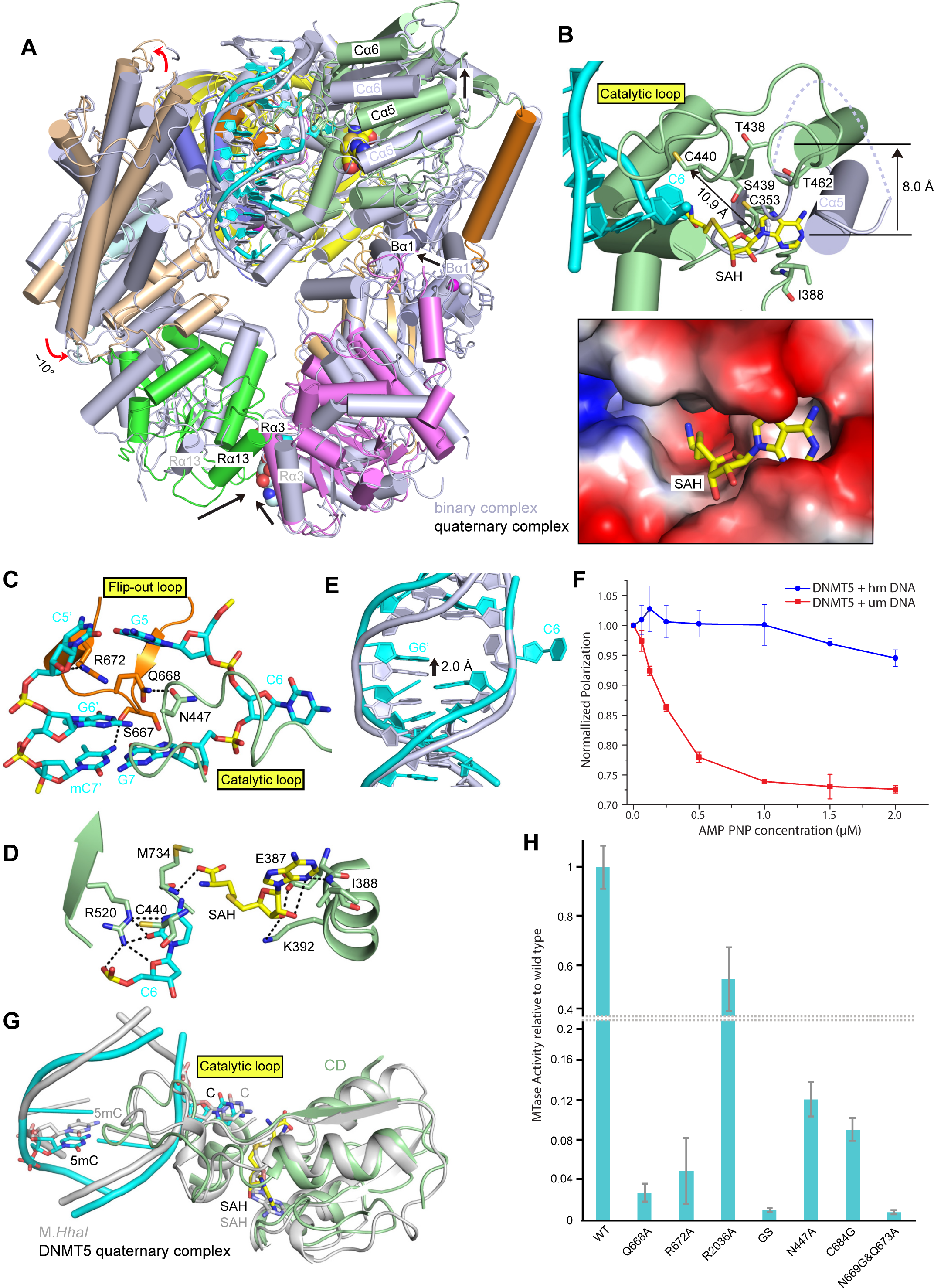
AMP-PNP Binding to ATPase Domain Activates the DNMT5 MTase Domain. (A) Conformational changes induced by AMP-PNP binding on proceeding from DNMT5 binary complex to the quaternary complex. (B) Conformational changes in the catalytic pocket on proceeding from DNMT5 binary complex (without bound SAH, in silver) to the quaternary complex (contains bound SAH, in color) (top) and electrostatic surface representation of the catalytic pocket in the quaternary complex (bottom). (C) Insertion of the catalytic loop into hm DNA and the complete base flipping of target cytosine. (D) Positioning of SAH within the MTase domain and its alignment relative to the target cytosine C6 and catalytic cysteine C440. (E) Overlay of the hmDNA from binary (in silver) and quaternary (in color) complexes. (F) FP assays of hmDNA and umDNA (FAM-labelled) preincubated DNMT5 with increasing amount of AMP-PNP. Error bars represent SD; n = 3. (G) Structural comparation of DNMT5 quaternary complex with M.*HhaI* ternary complex (PDB ID: 5MHT). (H) MTase activity of DNMT5 WT and mutants on hm DNA substrate. Error bars represent SD; n = 4. See also Figure S6.

The conformational changes associated with RecA1 propagate through the RecA1 insertions to the α/β region and upwards to the MTase domain. The MTase domain is lifted by the inward movements of α/β region such that the Cα5 helix following the catalytic loop is raised by 8 Å, allowing the catalytic loop to insert in the DNA (Figure 6B). Notably, the catalytic loop is visible along its entire length in the quaternary complex, in contrast to the majority of it being invisible due to flexibility in the binary complex. The side chain of N447 from the catalytic loop inserts into hmDNA from the minor groove direction, while the flip-out loop of the TRD approaches the DNA from the major groove direction (Figure 6C). Q668 from the flip-out loop, which recognized G6’ in the binary complex, now shifts to stabilize the catalytic loop by forming a hydrogen bond with N447 (Figure 6C). The target cytosine C6 flips completely out of the DNA duplex into the catalytic pocket of the MTase domain and is further stabilized by the side chain of R520 (Figure 6D). Importantly, the catalytic cysteine C440 within the catalytic loop also moves 10.9 Å from its position in the binary complex and locates near the target cytosine C6 (Figure 6B and Movie 2).

The insertion of the catalytic loop into hmDNA disrupts the hydrogen bonds stabilizing its small segment in the binary complex (Figure 4E) and opens the catalytic pocket for SAH binding (Figure 6B). SAH is bound in the proximity of the flipped-out cytosine and catalytic cysteine, with the adenosine base hydrogen-bonded to the main chain of I388, the sugar ring hydrogen bonded to the side chains of E387 and K392, and the carboxy tail stabilized by hydrogen bonding with M734 (Figure 6D). The binding affinity of SAH to DNMT5 is ∼15 μM in the presence of hmDNA and AMP-PNP (Figure 4G). The MTase domain, hmDNA and SAH in the DNMT5 quaternary complex superpose well (r.m.s.d. of 1.6 Å for 131 Cα atoms) with that of M*.HhaI* ternary complex (Figure 6G), suggestive of a potentially productive conformation.

Single turnover MTase experiments (enzyme excess) shows that AMP-PNP is able to induce MTase activity (Figure S6A). This activity is not due to AMP-PNP slow hydrolysis or contaminating ATP as hydrolysis product cannot be detected and even in the presence of hexokinase (Figure S6B). However, kinetic analysis showed a ∼1000-fold lower catalytic rate in the presence of AMP-PNP compared to ATP (Figure S7C). These findings indicate that nucleotide binding indeed places the enzyme is a conformation very close to the fully catalytic conformation, but hydrolysis is required for biologically meaningful activity.

### Nucleotide Binding Ensures Substrate Specificity

In addition, the hmDNA is lifted by 2 Å upon AMP-PNP binding (Figure 6E). This lift and the overall compact conformation in the quaternary complex would be a challenge for the non-specifically bound umDNA because of fewer contacts with DNMT5. We measured the fluorescence polarization of umDNA and hmDNA (FAM-labelled) preincubated DNMT5 with increasing amounts of AMP-PNP. Strikingly, the polarization of the DNMT5-umDNA complex decreased on addition of AMP-PNP, while that of the DNMT5-hmDNA complex is minimally affected (Figure 6F), demonstrating that non-specifically bound umDNA is dissociated from DNMT5 upon AMP-PNP binding. ITC experiments showed no detectable binding of SAH to umDNA pre-incubated DNMT5 in the presence of AMP-PNP (Figure 4G). Thus, the catalytic pocket of MTase domain remains closed with noncognate umDNA substrate, and AMP-PNP binding results in the release of the umDNA, which contrasts with opening the catalytic pocket for hmDNA substrate. These findings explain the lack MTase activity on umDNA in the presence of ATP (Figure 1B). We conclude that access of SAM to the catalytic pocket of DNMT5 MTase domain is controlled by its ATPase domain which licenses hmDNA for catalysis and ejects noncognate umDNA, which enables the system to achieve its remarkable specificity.

Finally, to test the relevance of the structural observations, we measured the MTase activity of DNMT5 mutants (Figure 6H). Alanine mutations of residues involved in 5mC recognition or partial flip-out of target cytosine, such as N669/Q673, Q668 and R672, exhibited predominant loss of MTase activity on hmDNA, emphasizing the importance of hmDNA recognition. Disruption of the interaction between HD and hmDNA backbone by the R2036A mutation reduced the catalytic activity by half relative to WT. The replaced GGS loop also shows low methylation activity. The relative MTase activity of N447A mutant, which displays no defect in ATPase activity (Figure 4C), is reduced by almost 10-fold, supporting the role for N447 insertion into the DNA duplex. The C684G mutant also exhibits a reduction in methylation activity. These data support the predictions of the structure data.

## DISCUSSION

DNMT5 is the only known ATP-dependent DNA cytosine-5 methyltransferase which displays exquisite specificity for hemimethylated DNA and, unexpectedly, mediates epigenome evolution over million-year timescales (Dumesic et al., 2020: Catania et al, 2020). Our structural and biochemical analysis reveals a stepwise allosteric cascade triggered by hmDNA binding that culminates in the completion of the construction of a MTase active site. Our studies reveal how communication between the ATPase and MTase domains occurs and how it is controlled by the DNA substrate and ATP. We discuss below how these communications occur at the molecular level to drive a hmDNA- and ATP-dependent allosteric cycle and how substrate recognition impinges on this cycle at multiple steps. Together, they provide a structural basis for understanding ATP-dependent epigenetic memory.

### hm DNA Recognition Triggers a Stepwise Allosteric Cascade in DNMT5

The ATPase domain of DNMT5 exhibits low basal activity in the absence of hmDNA substrate with its activity regulated by an autoinhibitory mechanism in which a pair of α helices in the ATPase domains are positioned to prevent RecA1 and RecA2 from forming a closed conformation (Figure 7A). hmDNA is recognized by DNMT5 through a ‘trigger mechanism’, like the first domino in a series falling, to reposition the side chain of N669 due to contacts by the methyl group of 5mC, which initiates the specific recognition of the orphan guanine by Q668 and subsequent conformational changes. Methylated CpG recognition induces the partial flip-out of the target cytosine from the DNA duplex where its unmethylated state is sensed by C684. The signal of hm DNA binding read by the MTase domain is communicated to the ATPase domain through release of the allosteric loop to contact with the RecA2-containing left segment of DNMT5, resulting in a profound rigid body rotation of the left segment on binary complex formation. The rotation stimulates the ATPase activity by aligning the two RecA-like domains for ATP binding and hydrolysis (Figure 7B).

**Figure 7.**
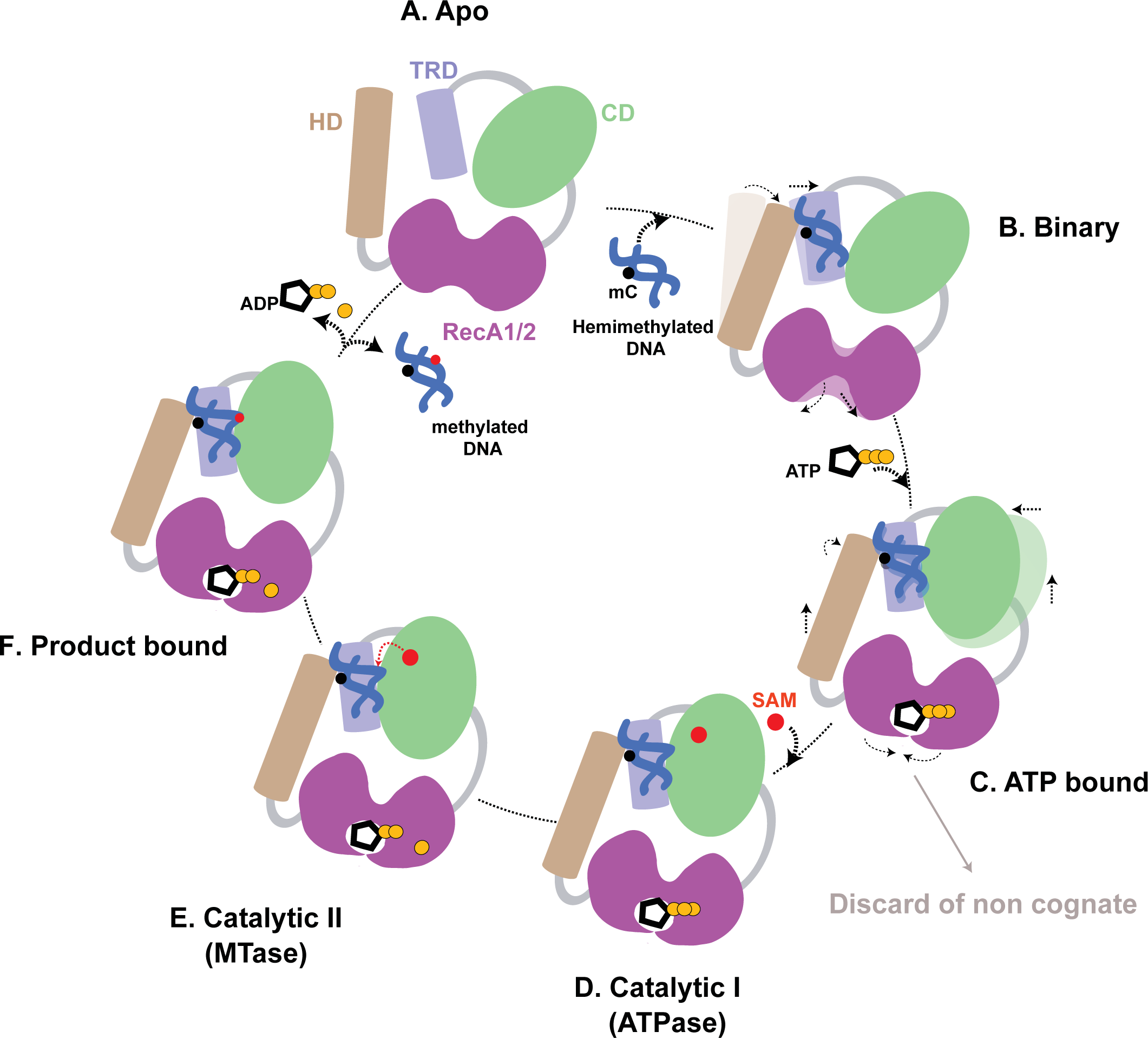
Model of DNMT5-mediated Methylation Cycle. (A) The ATPase domain adopts an autoinhibitory conformation that restricts DNMT5 with low basal activity in the apo state. (B) Specific recognition of hmDNA by the MTase domain stimulates the ATPase activity through a rigid body rotation to overcome the autoinhibitory conformation. (C) ATP binding to the ATPase domain remodels the MTase domain, opening the catalytic pocket for SAM binding on hmDNA substrate and licenses post-DNA-binding ejection on umDNA substrate. (D) Entry of SAM into the catalytic pocket. (E) ATP is hydrolyzed, fully activating the MTase to enable methyltransfer from SAM. (F) Product-bound state after catalysis.

Then, ATP-binding to the ATPase domain induces formation of a closed conformation, which is communicated to MTase domain mainly through the RecA1 insertions and the α/β region positioned within the right segment of DNMT5. The catalytic pocket of MTase domain is allosterically remodeled to promote catalytic loop insertion into the DNA duplex. The insertion opens the catalytic pocket for SAM entry, facilitates complete base-flipping into the active site and brings the catalytic cysteine into close proximity for methyl group transfer from SAM (Figure 7C). ATP binding also ejects noncognate umDNA to ensure substrate specificity. Subsequent SAM binding allows DNA methylation reaction to proceed (Figure 7D) and ATP hydrolysis to enable a fully catalytic conformation (Figure 7E).

### A Stable Partially Flipped Out State of Target Cytosine in DNMT5

We observed a novel stable partial flip-out state of target cytosine in the binary complex (Figure 2E) that has not been seen in other proteins and enzymes involved in writing, reading and erasing modified bases (Hong and Cheng, 2016; Klimasauskas *et al*., 1994). Methylated CpG dinucleotide recognition induces the target cytosine partially flip-out of the DNA duplex. DNA breathing (Chen and Prohofsky, 1995) may also play a role during partial base flipping. It appears that the partial flip-out of target cytosine does not need invasion of the catalytic loop into DNA duplex, given that the catalytic loop is invisible due to flexibility in the structure of the binary complex.

The partial flip-out state of target cytosine is critical for the eventual success of DNA methylation as mutations of residues involving in forming the intermediate state severely impair MTase activity (Figure 6H). The intermediate state may lower the free-energy barrier to achieve the completely flipped-out state. In addition, the partially flipped-out cytosine might constitute a second proofreading step following the initial 5mC recognition to discriminate hmDNA and fmDNA. It is worth noting that complete cytosine base flipping in DNMT5, not the partial flip-out state, needs ATP binding to the ATPase domain. Notably, a ^19^F NMR spectral study supports an ensemble of flipped-out forms equilibrating between a stacked form in the B-DNA and the completely flipped-out form (observed in the crystal structure of the M.*HhaI* ternary complex) (Klimasauskas et al., 1998). However, the flipped-out cytosine base is dynamic and lacks a clear preferred conformation in the above study, which is different from the stable partial flip-out state in DNMT5.

### The DNMT5 MTase Domain Requires Remodeling to be Active

Structure-based sequence alignment shows that the MTase domain of DNMT5 contains all conserved motifs of 5mC methyltransferases (Kumar et al., 1994; Lauster et al., 1989; Posfai et al., 1989), except some residue substitutions in motifs I-V that are responsible for SAM binding (Figures 4E, 4F, and S7). The conserved aromatic residues in motifs I and II of M.*HhaI* (F18 and W41) are replaced with small side chain residues (C453 and I388) in DNMT5. Additionally, hydrophobic residues (P80 and L100) in motifs IV and V of M.*HhaI* are changed to hydrophilic residues in DNMT5 (S439 and T462). By contrast to the tight binding of SAM to M.*HhaI* during purification (Kumar et al., 1992), these residue substitutions should weaken the binding of SAM to DNMT5 and make SAM binding regulatable. Importantly, the proline in the absolutely conserved catalytic cysteine-containing prolylcysteinyl (PC) dipeptide located in motif IV of M*.HhaI* (Kumar *et al*., 1992; Lauster *et al*., 1989; Posfai *et al*., 1989) is replaced by S439 in DNMT5. The *S. pombe* DNMT2 also has the serine substitution, which was thought to be functionally non-active as a result of the substitution (Pinarbasi et al., 1996). Afterwards, DNMT2 was shown to be a tRNA methyltransferase (Goll et al., 2006; Johannsson et al., 2018). Interestingly, these residue substitutions within DNMT5 generates a hydrogen-bond network stabilizing part of the catalytic loop that closes the catalytic pocket like a lid and prevents SAM from binding in apo-DNMT5 and binary complex. Additionally, hmDNA binds to DNMT5 in a position that is higher than that observed in the M.*HhaI* complex (Figure 4D), resulting in the DNMT5 catalytic loop unable to insert into the DNA duplex. These features mean that the MTase domain of DNMT5 must be remodeled to become active.

### A Novel Enzyme Remodeling Role of SNF2 ATPase Domain

SNF2 ATPases are best known for their role in chromatin remodeling enzymes (Flaus et al., 2006). Here we describe a SNF2 ATPase that remodels the catalytic pocket of the adjoined regulatable MTase domain to enable its remarkable substrate specificity. Nucleotide binding to the SNF2 ATPase domain allosterically opens the catalytic pocket in the MTase to allow SAM cofactor binding and triggers DNA modification specifically on the hmDNA substrate produced by DNA replication. Importantly, umDNA binding leads to a different outcome in which the pocket remains closed upon ATP binding but is instead ejected from the enzyme-substrate complex.

In this sense, the SNF2 ATPase domain of DNMT5 acts in a chaperone-like manner to complete folding and positioning of a catalytic domain into an active conformation with movement relative to the DNA. This active site remodeling allows SAM access to the catalytic pocket and the completion of base flipping into the active site. These findings help to explain the extraordinary specificity of DNMT5, which presumably enables it to mediate epigenetic evolution over geological timescales (Catania *et al*., 2020). As DNMT5 enzymes are found throughout the fungal and protist kingdoms, including in important photosynthetic picoeukaryotes (Huff and Zilberman, 2014; Ponger and Li, 2005), it is likely that the principles defined here apply to many organisms, which we predict will be found to harbor the ability to remember their past via DNA methylation, perhaps for long periods of time.

## ACKNOWLEDGEMENTS

We thank Geeta J. Narlikar and members of her laboratory, especially Emily Wong, for advice on the enzymological experiments. We thank Wei Xie, You Yu, Phillip A. Dumesic and Garrett M. Warren for helpful discussions and thank Stephen Long for use of the SpectraMax M5 microplate reader. D.J.P. is supported by funding from the Leukemia and Lymphoma Society, the Maloris Foundation and the Memorial Sloan-Kettering Cancer Center Core grant P30 CA008748. This work was also supported by NIH R01 GM071801 to H.D.M. H.D.M. is an Investigator of the Chan-Zuckerberg Biohub.

## AUTHOR CONTRIBUTIONS

J.W. undertook all the cryo-EM experiments with the help of C.W. and M.J.d.l.C., as well as the FP and ITC assays, under supervision of D.J.P. S.C. and H.D.M. designed and provided the insect cell constructs. S.C. and B.R. performed the ATPase and MTase activity assays under the guidance of H.D.M. J.W., D.J.P. and H.D.M. wrote the paper with input from S.C.

## STAR METHODS RESOURCE AVAILABILITY

### Lead contact and Materials availability

Further information and requests for resources and reagents should be directed to the lead contact Dinshaw J. Patel.

All reagents generated in this study are available from the Lead contact without restriction.

### Data and code availability

The cryo-EM density maps for apo-DNMT5, binary complex and quaternary complex have been deposited in the EM Database under accession codes: EMD-24292, EMD-24294, EMD-24295 and the coordinates for the structures have been deposited in Protein Data Bank under accession codes: 7R76, 7R77, 7R78.

## METHODS DETAILS

### Protein Expression and Purification

The gene encoding *C. neoformans* DNMT5 (residue 58-2377) was codon optimized for insect cell expression and inserted into pFastBac HT B vector (Life Technologies) with an N-terminal 6×His tag. Recombinant bacmid was generated in DH10Bac *E. coli* (Life Technologies) and then transfected into adherent *Spodoptera frugiperda* Sf9 cells (Life Technologies) for recombinant baculovirus packing. Supernatant containing the baculovirus was collected for reinfection and amplification. Suspension *Trichoplusia ni* High5 cells (ThermoFisher Scientific) were infected by high titer baculovirus and collected after 72 h infection. The recombinant DNMT5 was purified by 5 ml HisTrap Fastflow column (GE Healthcare) and dialyzed to low salt buffer (20 mM Tris, pH 8.0, 150 mM NaCl, 5 mM β-mercaptoethanol). The supernatant was further purified by HiTrap Q FF column and Superdex 200 increase 10/300 column (GE Healthcare). The pooled fractions were used for subsequent analysis or flash-frozen in liquid nitrogen.

DNMT5 mutations were generated using the Q5 site-directed mutagenesis kit (New England Biolabs) and verified by sequencing. All DNMT5 mutants were expressed and purified following the same protocol as the wild-type protein.

### Cryo-EM Sample Preparation

For the DNMT5 apo form and binary complex, the purified DNMT5 was mixed with the 36-bp hm DNA, AMP-PNP, SAH at the molar ratio of 1:5:10:10 in the presence of 2 mM MgCl_2_ and incubated on ice for 1 hour. The mixture was further purified by gel filtration on the Superdex 200 increase 10/300 column (GE Healthcare) in buffer 20 mM Tris, pH 8.0, 150 mM NaCl, 2 mM DTT. The peak fractions containing DNMT5 complexes were pooled and concentrated at around 0.7 µM (OD_280_∼0.4, OD_260_/OD_280_=0.97) for cryo-EM grid preparation.

For the quaternary complex, the purified DNMT5 was diluted to final concentration of ∼0.2 mg/ml and incubated with the hm DNA, AMP-PNP and SAH at the molar ratio of 1:2:10:10 in buffer 20 mM Tris, pH 8.0, 150 mM NaCl, 2 mM DTT, 2 mM MgCl_2_ for 1 hour on ice. The mixture was subject to grid preparation directly without further purification.

The pseudo ternary complex was prepared as for the quaternary complex except that SAH was changed to SAM and AMP-PNP was absent.

### Cryo-EM Sample Data Acquisition

3.5 μl peak fractions or incubated complexes were applied onto glow discharged UltrAuFoil 300 μ mesh R1.2/1.3 grids (Quantifoil) at ∼4°C. Grids were blotted for 1.5 s at 100% humidity and flash frozen in liquid ethane using a Vitrobot Mark IV (FEI/Thermo Fisher Scientific). Images were collected on a Titan Krios G2 (FEI/Thermo Fisher Scientific) transmission electron microscope operating at 300 kV with a K3 direct detector (Gatan) using a 1.064 Å pixel size at the Memorial Sloan Kettering Cancer Center’s Richard Rifkind Center for cryo-EM. The defocus range was set from -1.0 to -2.5μm. Movies were recorded in super-resolution mode at an electron dose rate of 20 e^-^/ pixel/s with a total exposure time of 3 s and intermediate frames were recorded every 0.075 s for a total of 40 frames.

### Image Processing

For the DNMT5 apo form and binary complex datasets, stacks were motion corrected and 2× Fourier-cropped from a super-resolution pixel size of 0.532 Å using MotionCor2 (Zheng et al., 2017). Contrast transfer function parameters were estimated by CTFFIND-4 (Rohou and Grigorieff, 2015). Other steps of cryo-EM data processing were performed by RELION-3 (Zivanov et al., 2018). Particles were auto-picked without reference and extracted at a 3-fold binned pixel size of 3.192 Å. After two rounds of reference-free 2D classification, 715,061 particles were selected for 3D classification using an initial model as reference generated by RELION. 325,656 particles from the best two classes were refined, re-extracted to pixel size 1.064 Å and used for consensus 3D refinement. The resulting map had well-resolved density in the right segment and rather weak density in the left segment, indicating conformational heterogeneity. We tried to perform masked 3D classification on the left segment with the signal of the right segment subtracted from each raw particle image (Bai et al., 2015). The 325,656 density-subtracted particles were subject to 3D classification, while keeping all orientations fixed at the values determined in the consensus 3D refinement map. Classification into 8 classes yielded 2 major classes that showed good density (Figure S1A). The original and non-subtracted particles from the 2 good classes were used to perform 3D refinement separately. Postprocessing in RELION-3 (using pixel size 1.064 Å) yielded the final reconstructions at 3.0 Å for DNMT5 binary complex and 3.2 Å for apo-DNMT5.

The dataset of the quaternary complex was processed by a similar procedure as above. Briefly, 2,439,273 particles were auto-picked from 2,608 movies and 1,700,803 particles were selected for 3D classification after two rounds of reference-free 2D classification. After three rounds of 3D classification, 23,621 particles from the best class were subject to 3D refinement along with CTF refinement and particle polishing. The final reconstruction yielded the structure of the quaternary complex with an overall resolution of 3.5 Å.

All resolutions were estimated using RELION post-processing by applying a soft mask around the protein density and the Fourier shell correlation (FSC)=0.143 criterion (Scheres and Chen, 2012). The local resolution was calculated with RELION-3.

### Atomic Model Building and Refinement

The atomic model of the binary complex was *de novo* built manually in Coot (Emsley et al., 2010) based on the bulky amino acid side chains in the primary protein sequence. The apo-DNMT5 and quaternary complex models were built by docking the right and left segments of the binary complex model separately into the cryo-EM maps using UCSF Chimera (Pettersen et al., 2004) and then rebuilt and confirmed by bulky side chains in Coot. These models were refined by real-space refinement in Phenix (Adams et al., 2010) by applying geometric and secondary structure restraints. Structure figures were prepared with Pymol (pymol.org), UCSF Chimera and Chimera X (Goddard et al., 2018).

### Fluorescence Polarization (FP) Assay

FP assays were performed in black flat-bottom 384-well plates (Greiner Bio-one) with the final assay volume of 20 µl per well. DNMT5 or DNMT5-DNA mixture were serially diluted 2-fold in a buffer containing 20 mM Tris, pH 8.0, 50 mM NaCl, 2 mM MgCl_2_, 2 mM DTT (unless otherwise indicated).

For DNMT5 binding to DNA assays, 6-carboxyfluorescein (FAM)-labeled double-stranded DNA (dsDNA) was generated by annealing equimolar amounts of sense and antisense oligonucleotides (IDT) using a PCR machine. Serially diluted DNMT5 were incubated with a final 50 nM concentration of FAM-labelled dsDNA at room temperature (RT) for 20 min. Polarizations were measured on a Tecan infinite m1000 microplate reader at an excitation wavelength of 470 nm and an emission wavelength of 520 nm. The polarization values measured from control wells containing only FAM-dsDNA (in absence of protein) were considered as background polarization signal and subtracted from polarization values measured from experiment wells containing protein. Scatter plots of polarizations versus DNMT5 concentration were plotted using Origin software (OriginLab) and dissociation constants (K_d_) were fitted with a OneSiteBind model.

For AMP-PNP competition FP assay, 1 µM DNMT5 was incubated with 100 nM FAM-labelled hm or um DNA on ice for 1 hour. The DNMT5-DNA mixtures were titrated with 2-fold serially diluted AMP-PNP at 1:1 volume ratio. Following a 20 min incubation period at RT, the polarizations were measured as above. The polarization values measured from control wells containing only DNMT5-dsDNA (in absence of AMP-PNP but same volume of buffer added) were normalized as 1. The normalized polarizations versus AMP-PNP concentration were plotted using Origin software.

For the ATP binding assays, DNMT5 was incubated with 4-fold molar excess hm DNA on ice for 1 hour and then 2-fold serially diluted. In the group without hm DNA, the same volume of buffer was added. 0.2 µM Fluorescent ATP analog 2’(3’)-O-(N-methyl-anthraniloyl)-AMP-PNP (mantAMP-PNP) (Biorbyt) was mixed with serially diluted DNMT5 or DNMT5-hm DNA and incubated for 20 min at RT. Fluorescence polarization was measured on a SpectraMax M5 microplate reader (Molecular Devices LLC) with an excitation wavelength of 355 nm and an emission wavelength of 448 nm.

### Isothermal Titration Calorimetry (ITC)

Purified DNMT5 was incubated with 4-fold molar excess hm or um DNA overnight on ice and 5-fold molar excess AMP-PNP or equivalent volume buffer were added to incubate on ice for another 1 hour. The final concentration of DNMT5 is ∼10 µM. SAH was dissolved to 200 µM in the same buffer as DNMT5 protein containing 20 mM Tris, pH 8.0, 50 mM NaCl, 2 mM MgCl_2_, 5mM β-mercaptoethanol. The titrations were performed on an iTC200 MicroCalorimeter (Malvern Panalytical) at 20°C with 300 rpm stirring speed. Each ITC titration consisted of 20 successive injections with 0.4 µl for the first injection and 2.0 µl for the rest. The resultant ITC curves were processed using Origin software and fitted according to ‘One Set of Sites’ model with the stoichiometry (n value) fixed at 1 for valid curve fitting (Turnbull and Daranas, 2003).

### ATPase Assay

ATPase activity was assessed as described in (Catania *et al*., 2020). In brief, 30-60 nM of recombinant DNMT5 was incubated in ATPase reaction buffer (50 mM HEPES-KOH pH 7.9, 75 mM KCl, 5% Glycerol, 2 mM DTT, 2mM phosphoenolpyruvate (Sigma), 0.18 mM NADH (Sigma), 0.5 mM ATP, 5 mM MgCl_2_, 10 U/ml pyruvate kinase (Sigma), 10 U/ml lactate dehydrogenase (Millipore). DNA substrates were added to a final concentration of 5 µM and the reactions performed at 25°C in a 384 well non-stick clear bottom plates (Corning 3766). The absorbance at 340 nm and 420 nm were measured using a Tecan Spark 10M. The difference between 340 nm and 420 nm was plotted versus time and the initial rate was calculated in the linear portion of the curve. The rate values were normalized over the concentration of DNMT5 used in the assay.

To measure ATPase activity in single turnover experiments (Figure S6), ADP produced from the reaction was measured using ADP-Glo Kinase assay (Promega) following the manual instructions. DNMT5 (1 µM) was incubated for 4 h in methyltransferase buffer + glucose (50 mM Tris-HCl pH 8, 25 mM NaCl, 10% Glycerol, 2mM DTT, 1 mM Glucose, 5 mM MgCl_2_) in the presence of 0.5 mM ATP or AMP-PNP in a total volume of 5µl. 5µl of Reagent solution was added and incubated for 40 min at RT followed by addition of 10µl of Detection solution and incubation for 60 min. The luminescence was detected using a Synergy H1 plate reader (BioTek). For some of the samples, a pre-incubation step with Hexokinase (36U/ml, Sigma H-6380) was performed for 1h at RT before addition of DNMT5.

### DNA Methyltransferase Assay

The DNA methyltransferase assays were carried out as described in (Roth and Jeltsch, 2000) with some modifications. DNMT5 (30 nM) was incubated at RT in methyltransferase buffer (50 mM Tris-HCl pH 8, 25 mM NaCl, 10% Glycerol, 2mM DTT, 1 mM ATP, 1 mM MgCl_2_) in the presence of 5 µM biotinylated DNA substrate and 4 µM ^3^H-SAM (PerkinElmer). 5 µl of reaction were removed at indicated time points and quenched in binding buffer (5 mM Tris-HCl pH 7.5, 0.5 mM EDTA, 1 M NaCl) containing 10 mM SAM in 10 mM H_2_SO_4_. At the end of the time course, the reactions were transferred to 96 well streptavidin-coated plates (Sigma-S2577) and incubated for 30 min at RT. The plates were washed 3 times in 200 µl of binding buffer and 3x 200 µl Digestion Buffer (50 mM Tris-HCl pH8, 5 mM MgCl_2_). To release the radioactivity incorporated in the DNA, the plates were incubated with 25U Benzonase Nuclease (Sigma) in 100 µl Digestion buffer for 1 h at RT. The reactions were then transferred in 3 ml of scintillation liquid (Ultima Gold-PerkinElmer) and the radioactivity detected on an LS 6500 scintillation counter (Beckman Coulter). Background signals (reaction without enzyme) were generally between 3 to 7 CPM. The initial rate values were calculated in the linear portion of the curve and normalized over the concentration of DNMT5. For the End point assays, the reactions were performed using the same protocol described above but in the presence of 100 nM DNMT5 and for 4 h.

Single turnover experiments were performed as described above but incubating 1 µM DNMT5 with 100 nM biotinylated hm DNA (possessing a single mCpG site) at RT and using 1 mM ATP or AMP-PNP.

---

**Figure S1.**
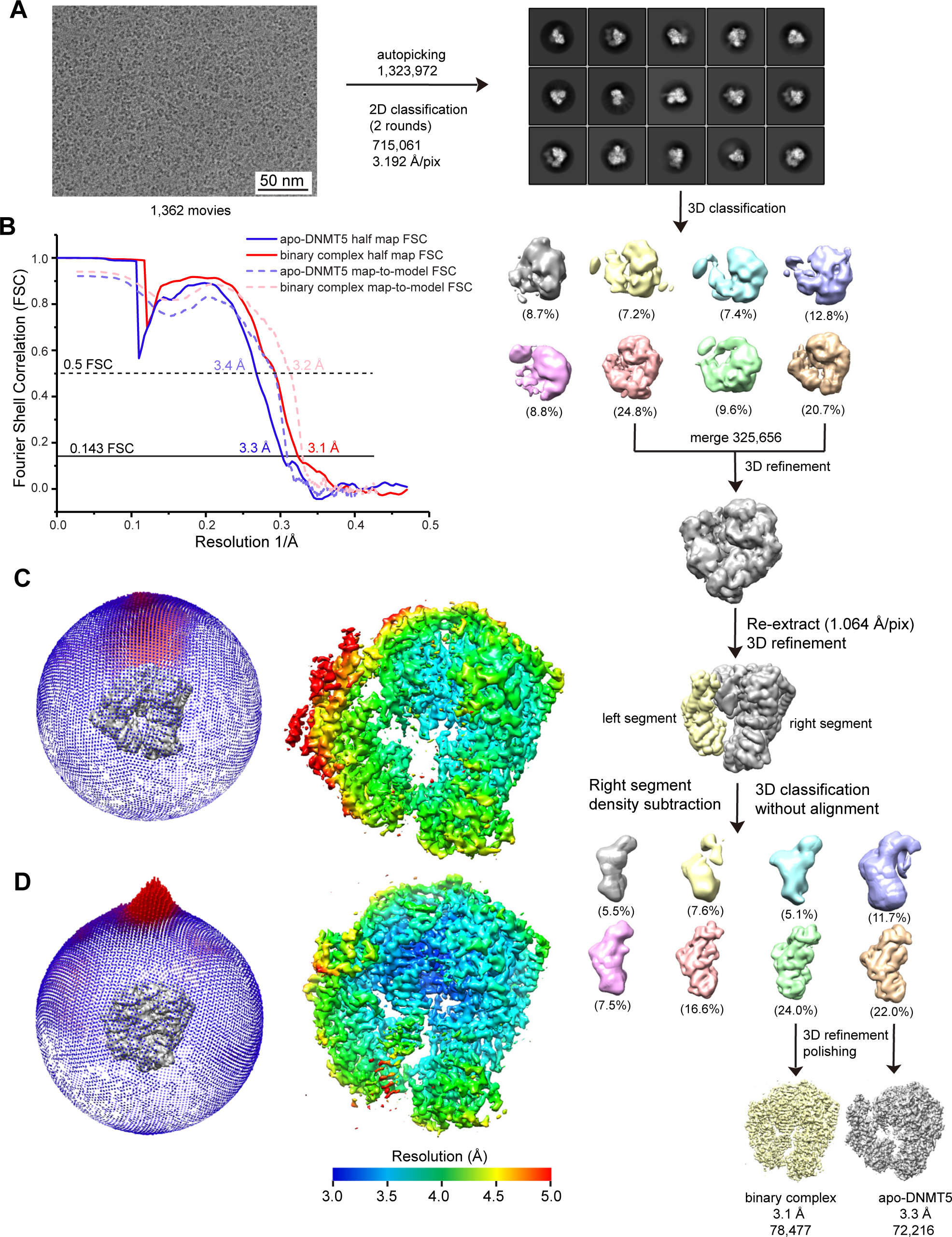
Cryo-EM reconstructions of DNMT5 and its binary complex with hmDNA. (A) Flow chart of image processing of DNMT5 and its binary complex. (B) Fourier Shell Correlation (FSC) curves of half map and map-to-model for apo-DNMT5 and its binary complex. (C and D) Euler angle distributions (left panel) and final 3D reconstructed maps colored according to local resolution (right panel) of DNMT5 (C) and its binary complex (D).

**Figure S2.**
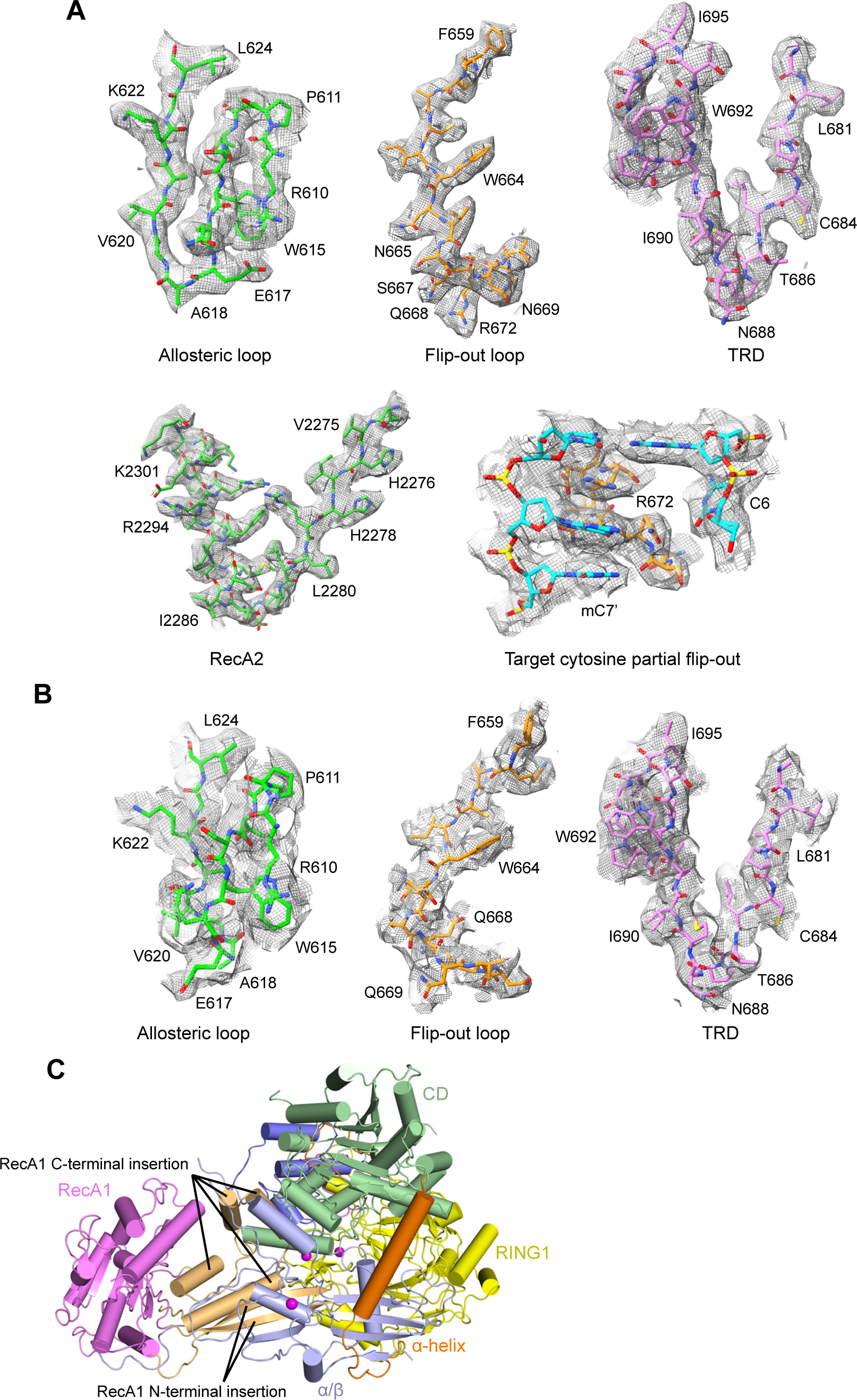
Cryo-EM Density Maps of DNMT5 and its Binary Complex and Folding Elements in the Structure of apo-DNMT5. (A and B) Density maps for indicated regions in DNMT5 binary complex (A) and apo-DNMT5 (B) maps at 0.015 contour level. (C) Side view of apo-DNMT5 structure showing the N- and C-terminal insertions of RecA1.

**Figure S3.**
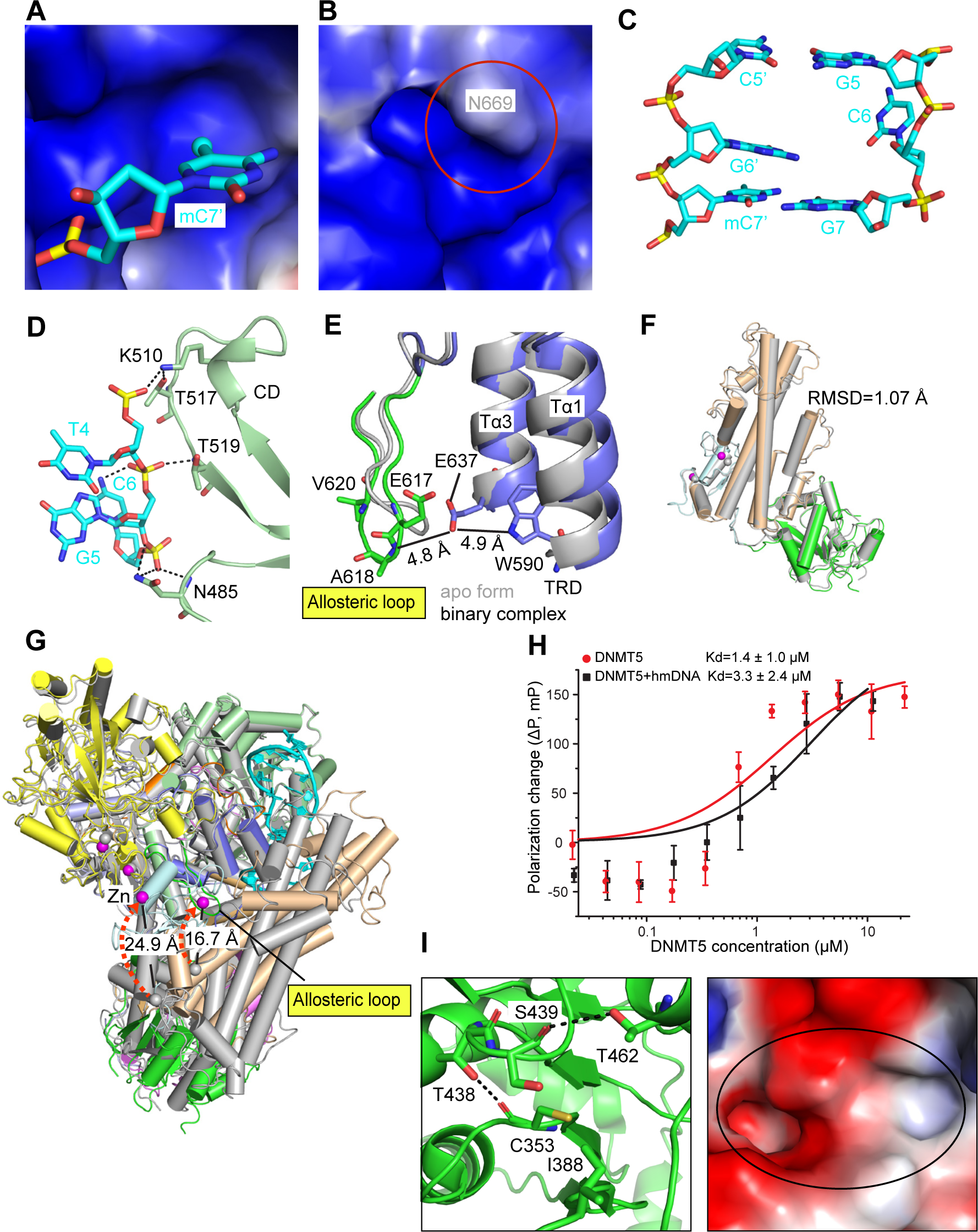
Interactions between DNMT5 and hmDNA. (A and B) Electrostatic surface representation of the basic pocket accommodating 5mC in the binary complex (A) and apo-DNMT5 structures (B). (C) The partial flip-out of target cytosine in the pseudo-ternary complex structure. (D) Interactions of DNMT5 with the phosphate backbone of hmDNA unmethylated strand. (E) hmDNA binding releases the allosteric loop by disrupting the hydrogen-bond network. (F) Overlay of the left segment of apo-DNMT5 (in silver) and its binary complex with hmDNA (in color). (G) The two zinc atoms in RING2 move by 24.9 Å anf 16.7 Å, respectively on proceeding from apo-DNMT5 (in silver) to its binary complex with hmDNA (in color). (H) Fluorescence polarization assay of DNMT5 alone and hmDNA preincubated DNMT5 with increasing amount of added mant-AMP-PNP. Error bars represent; SD = 3 (I) Ribbon (left) and electrostatic surface (right) representation of the catalytic pocket in apo-DNMT5 structure.

**Figure S4.**
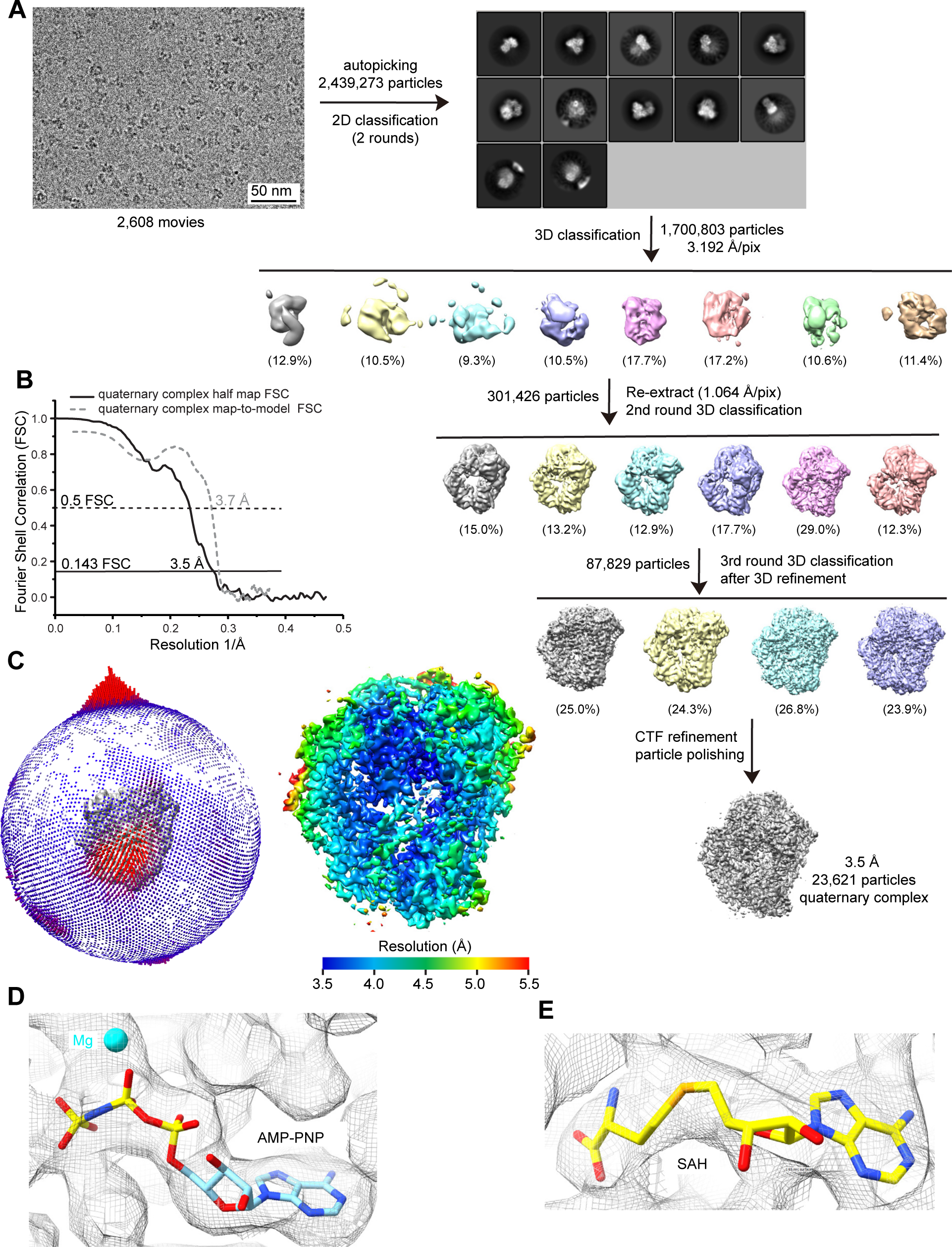
Cryo-EM reconstruction of DNMT5 quaternary complex with hmDNA, AMP-PNP and SAH. (A) Flow chart of image processing of DNMT5 quaternary complex structure. (B) Fourier Shell Correlation (FSC) curves of half map and map-to-model for the quaternary complex structure. (C) Euler angle distribution (left panel) and final 3D reconstructed map of the quaternary complex colored according to local resolution (right panel) of DNMT5 quaternary complex. (D and E) Density maps for AMP-PNP bound in the ATP-binding pocket (D) and for SAH bound in the catalytic pocket (E) in the quaternary complex map at 0.015 contour level.

**Figure S5.**
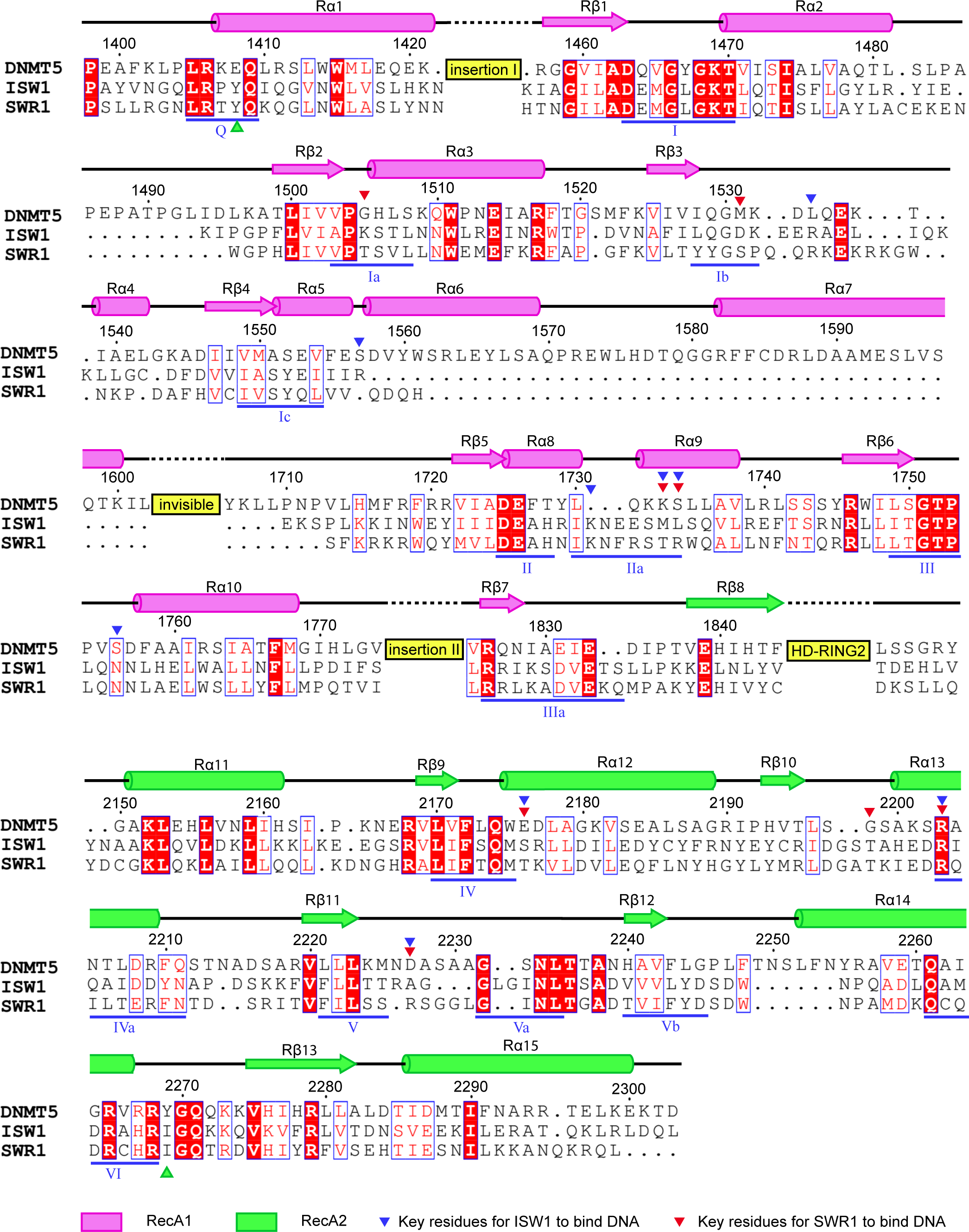
Structure-based Sequence Alignment of ATPase Domains from DNMT5, ISW1 and SWR1. Blue triangles show key residues of ISW1 for interaction with nucleosome DNA, while red triangles show key residues of SWR1 for interaction with nucleosome DNA. The PDB IDs are 6IRO (ISW1) and 6GEN (SWR1).

**Figure S6.**
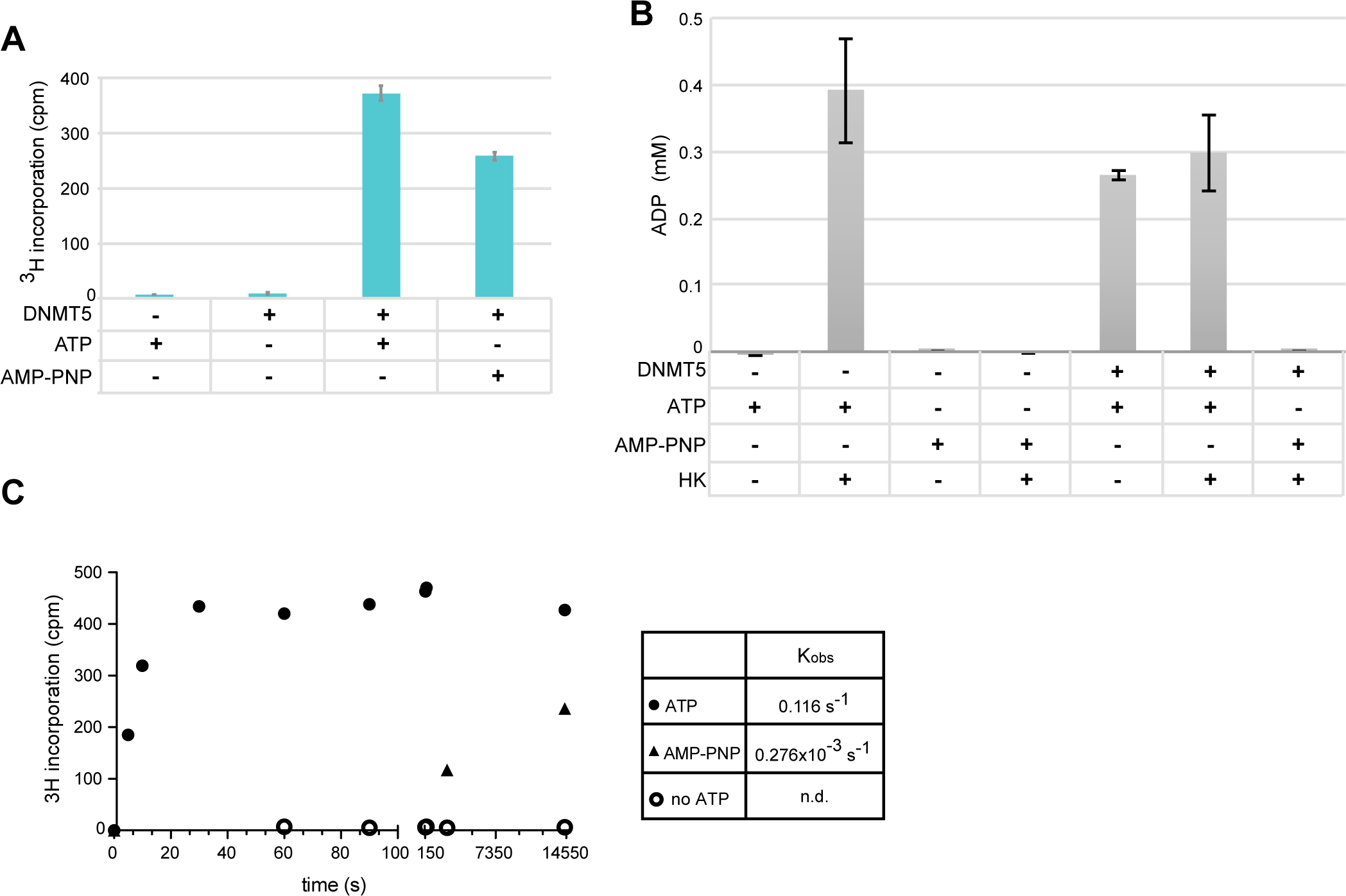
AMP-PNP induces slow but detectable MTase activity. (A) Single turnover end point MTase assay using 1 µM DNMT5 incubated with 100 nM hmDNA in the presence of 1 mM ATP or AMP-PNP for 4h at RT. (B) ATPase assays measure ADP produced by the ATPase activity of the enzymes using the same condition as in panel (A). Where indicated, samples were treated with the ATPhase hexokinase in the presence of 1 mM Glucose before the addition of DNMT5. (C) Time course of the MTase assays with same condition as in panel A and in the presence of ATP and AMP-PNP or no nucleotide (no ATP). Data in A-C are represented as mean ± s.d. from n=6 independent samples.

**Figure S7.**
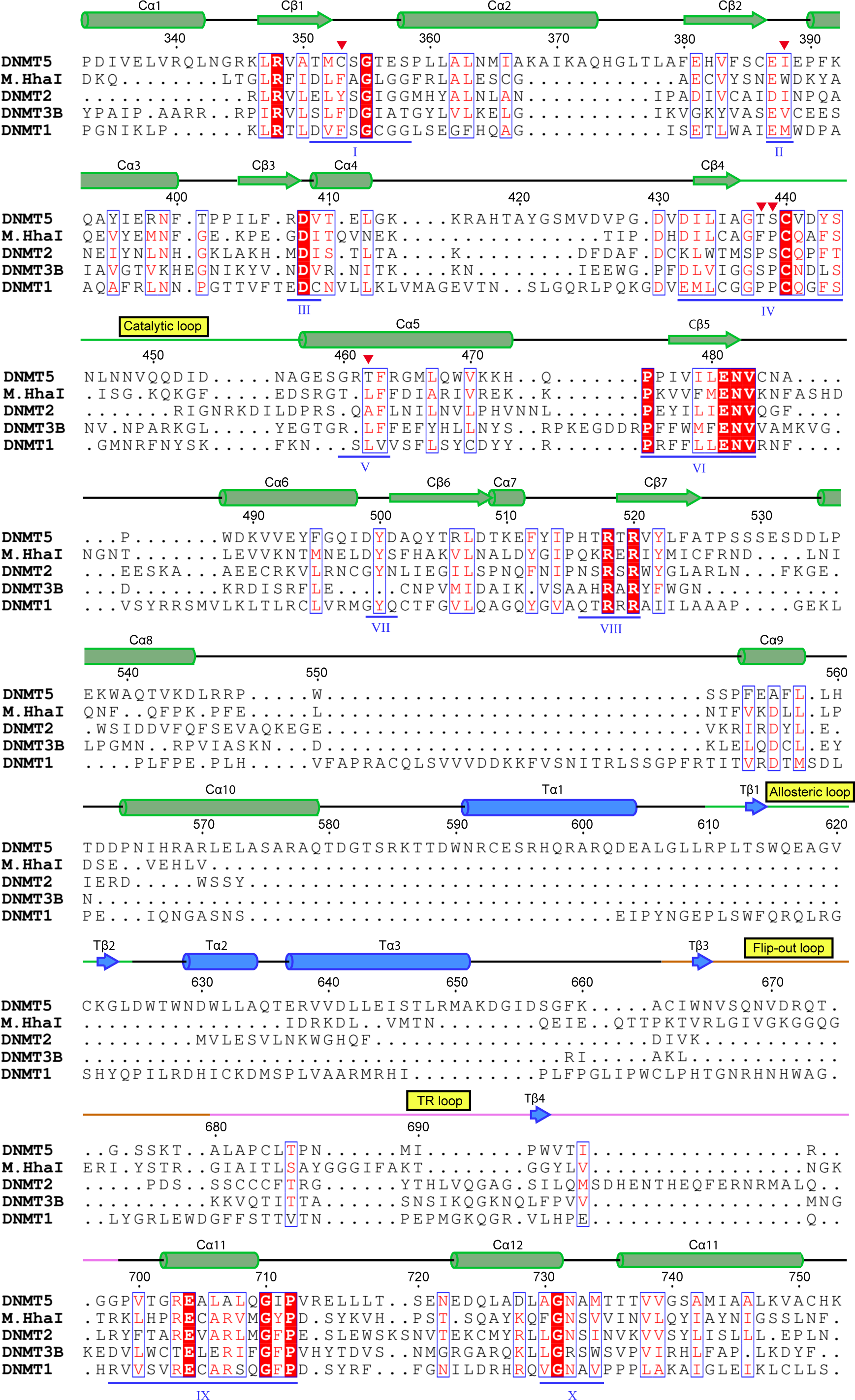
Structure-based Sequence Alignment of MTase Domains from DNMT5, M.HhaI, DNMT2, DNMT3B and DNMT1. Red triangles show residue substitutions in the catalytic pocket for SAM binding. The PDB IDs are 3EEO (M.HhaI), 6FDF (DNMT2), 6U8W (DNMT3B) and 6W8W (DNMT1).

